# Increasing Muscle Hypertrophy with a Natural Product Designed to Inhibit SIRT1

**DOI:** 10.1101/2022.06.27.497838

**Authors:** Suraj Pathak, Marita Wallace, Sonia Athalye, Simon Schenk, Henning T. Langer, Keith Baar

## Abstract

Muscle mass and strength are predictors of longevity. We have previously identified a series of molecular brakes that slow muscle growth in response to stress. One potential stress that we hypothesized would limit muscle growth is caloric stress through the activation of SIRT1. We therefore identified natural product inhibitors of SIRT1 and tested their effects on load-induced increases in muscle fiber cross-sectional area (fCSA) using an incomplete factorial design. Supplying varying amounts of three natural products for the full two-week period of overload resulted in increases in fCSA that varied from −2 to 113%. Using these data, we produced a model that predicted the optimal combination and concentration of each natural product and validated this model in a separate cohort of animals. Following two weeks of overload, fCSA in the optimal group increased 62%, whereas in the placebo fCSA increased only 3%. The greater increase in fCSA was not the result of an increase in ribosomal mass. In fact, the optimal group showed significantly less of the 5⍰ external transcribed spacer, a marker of 47S ribosomal RNA synthesis, and a trend for decreased total RNA. In spite of the lower ribosome mass, the increase in protein synthesis was similar, suggesting that the natural product cocktail may be increasing ribosomal efficiency rather than capacity. These data suggest that inhibition of SIRT1, together with exercise, may be useful in increasing muscle fCSA.

## Introduction

Muscle mass and strength are important aspects of human health, since the rate of mortality of individuals correlates with both low muscle mass (26) and strength (23, 24). Low muscle mass and function also limits post-surgery recovery and mobility (12), and increases the impact or risk of diseases such as diabetes, cardiovascular disease, and cancer (2). Thus, improving muscle mass and strength is a vital part of life- and health-span (8).

Muscle mass and strength are also important components of human aesthetics and performance. Indeed, each year, tens of billions of dollars are spent on supplements that are purported to result in increases in muscle mass and strength (21). Many of these products are of dubious scientific value, since wide scale screens for products that result in *bona fide* improvements in muscle mass and strength are rare. One clinically validated way to increase muscle mass gains as a result of training is to combine strength training with protein supplementation (3). However, very few scientifically validated nutraceutical approaches to increase muscle mass in response to training have been reported.

Sirtuin1 (SIRT1) is an NAD^+^-dependent deacetylase that is activated in muscle in response to changes in cellular energy flux. Metabolic stress during calorie restriction (6) and endurance exercise (3) are known to directly activate SIRT1. Since calorie restriction and endurance exercise are thought to slow muscle growth (1,7), we hypothesized that SIRT1 inhibits muscle growth. Protein acetylation has also been linked with muscle growth. The ribosomal S6 protein kinase (S6K1), whose phosphorylation and activity has previously been shown to be associated with increased muscle protein synthesis and muscle hypertrophy (1), can be acetylated by the acetyl transferase p300 and deacetylated by SIRT1 (15). Beyond S6K1, almost every protein within the ribosome has at least one acetylated lysine (4). Given that lysine acetylation can regulate numerous aspects of cellular homeostasis beyond its well-described role in epigenetics and gene transcription, it is possible that acetylation contributes to the regulation of protein synthesis. Since loading and nutrition result in transient increases in myofibrillar protein synthesis (7, 8), which is thought to play an important role in muscle growth (5), here we sought to determine whether increasing acetylation within muscle could augment the increase in muscle fiber cross-sectional area in response to a hypertrophic stimulus.

Our approach to addressing this objective to was to increase protein acetylation by reducing SIRT1 deacetylase activity. For this, we first sought to identify natural products that could inhibit SIRT1 deacetylase activity. Second, we sought to determine whether these natural products could augment muscle hypertrophy when combined in the optimal manner. Overall, our hypothesis was that a novel nutritional supplement that inhibits SIRT1 deacetylase activity could augment the increase in muscle fiber cross-sectional area in response to an overload stimulus.

## Materials and Methods

### SIRT1 inhibitor screen

The NatProd Collection library (MicroSource Discovery Systems, Inc. Gaylordsville, CT) in ten source plates was screened at two doses (5 μM and 50 μM final in the reaction mixture, duplicate for each dose). The inhibitory activity of the compounds was assessed in duplicate against 2ng/µL purified human SIRT1 using 3µM p53 as a substrate. The assay was a mobility shift assay based on charge differences before and after electrophoretic separation of product from fluorescently labeled substrate read using a Caliper EZ Reader (Perkin Elmer, Boston, MA). All reactions took place in the presence of excess nicotinamide adenine dinucleotide and the known SIRT1 inhibitor suramin was used as a control.

### Box-Behnken model generation

Three of the inhibitors identified in the natural product screen were selected based on their previous use in humans and complementary chemical structures. These inhibitors were then used to create an incomplete multifactorial design Box-Behnken model using Design-Expert® software. The three-factor design with one central point required 12 animals. Five more animals received the central dose of all three natural products to determine the biological variability of the intervention.

### Synergist ablation

All animal procedures were approved by the Institutional Animal Care and Use Committee at the University of California, Davis. Eighteen rats were used for both the DOE and validation experiments. The animals were anesthetized using 2.5% isoflurane, the distal hindlimb shaved and prepared for aseptic surgery. The whole soleus and bottom two thirds of the gastrocnemius were removed at the Achilles tendon, leaving the plantaris (PLN) muscle and innervating nerve intact. The overlying fascia and skin were sutured closed, and the animals were moved to a temperature regulated area for recovery. The left leg served as a contralateral control. Animals were monitored daily to ensure they returned to normal activity and did not suffer any stress from the procedure. Animals were orally gavaged on a daily basis (just prior to lights out), in accordance with their respective treatment group.

### Muscle Collection

Following the 14th day of treatment, animals were anesthetized, and the overloaded and contralateral PLN muscles, hearts, and livers were collected. Upon removal, PLN muscles were trimmed conservatively, weighed and then pinned at resting length on cork, snap-frozen in liquid nitrogen cooled isopentane and stored at −80°C until further processing.

### Histology

PLN muscles were blocked in a cryostat on corks using Tissue Tek O.C.T. Compound (Sakura), and 10µm sections were mounted onto slides for CSA quantification. Slides were prepped for histological analysis by being blocked in 5% normal goat serum (NGS) in PBST w/1% tween, and incubated in primary antibodies for type I, IIa, IIb fibers, and/or laminin over night at 4°C. The next day, slides were washed with PBST w/0.1% Tween and incubated in HRP conjugated secondary antibodies for 60 minutes, washed again and mounted using Prolong Gold (no Dapi). Four random images were taken of each respective muscle section for CSA quantification using Fiji.

### Validation experiment

Following synergist ablation, animals were randomly assigned to one of four treatment groups, control (n=3), Least (n=5), Moderate (n=5), and Optimal (n=5). Least, moderate, and optimal refer to the best (optimal) and worst (least) combination of the natural products relative to the predicted effect on muscle fiber CSA. The Moderate group was predicted to have an effect of fiber CSA between that of the Optimal and Least groups. The control group received phosphate buffered saline (PBS), whereas the least, moderate, and optimal groups received different combinations and concentrations of the three SIRT1 inhibitors as a single cocktail dissolved in PBS (Table 2). All treatments were administered via oral gavage just prior to lights out for 14 days; the first gavage was given at 5pm on the day the surgery took place.

### mRNA Isolation, reverse transcription, and qPCR

Following blocking, PLN muscles were powdered using a hammer and pestle. Total RNA was extracted from weighed powdered muscle tissue using RNAzol in accordance with the manufacturer’s protocol. RNA was quantified using Biotek Epoch Microplate Reader via absorbance (Biotek,Winooski, VT), and the RNA/mg muscle was calculated by dividing total RNA in the sample by the mass of muscle powder used. 1.5µg of total RNA was then converted into cDNA using MultiScribe reverse transcriptase and oligo (DT) primers. cDNA was diluted 1:10 before conducting qPCR. qPCR was performed using CFX384 Touch Real-Time PCR Detection System (Bio-Rad, Hercules, CA), along with Quantified Mastermix and Bio-Rad Sybr Green Mix solution and Bio-Rad 384 well PCR plates. PCR reactions were performed in accordance with the manufacturer’s instructions with the following primers: rITS-1 (fwd-TCCGTTTTCTCGCTCTTCCC-; rev-CCGGAGAGATCACGTACCAC-), r5E1TS (fwd-ACGCACGCCTTCCCAGAGG-; rev-CGCGTCTCGCCTGGTCTCTTG-). Gene expression was calculated using a delta delta threshold cycle method (19) and GAPDH (fwd-TGGAAAGCTGTGGCGTGAT-; rev-TGCTTCACCACCTTCTTGAT-) was used as the housekeeping gene. The absolute C_T_ of GAPDH was unchanged by either overload or treatment.

### Tissue homogenization and western blotting

Two scoops of muscle powder were incubated in 250 µL of sucrose lysis buffer [1M Tris, pH 7.5, 1M sucrose, 1mM EDTA, 1mM EGTA, 1% Triton X-100, and protease inhibitor complex]. The solution was set on a shaker for 60 minutes at 4°C, spun down at 8,000g for 10 minutes, supernatants were transferred to new Eppendorf tubes and protein concentrations were then determined using a DC protein assay (Bio-Rad, Hercules, CA). 750 µg of protein was diluted in 4X Laemmli sample buffer (LSB) and boiled for 5 minutes. 10µL of protein sample was loaded onto a Criterion TGX Stain-Free Precast Gel and run for 45 minutes at a constant voltage of 200V. Proteins were then transferred to an Immobilon-P PVDF membrane, after it was activated in methanol and normalized in transfer buffer, at a constant voltage of 100V for 60 minutes. Membranes were blocked in 1% Fish Skin Gelatin (FSG) in TBST (Tris-buffered saline w/ 0.1% Tween) and incubated overnight at 4°C with the appropriate primary antibody diluted in either TBST or 1% FSG at 1:1,000. The next day, membranes were washed three times with TBST for 5 minutes, and successively incubated at room temperature with peroxidase-conjugated secondary antibodies in a 0.5% Nonfat Milk TBST solution at 1:5,000. Bound antibodies were detected using a chemiluminescence HRP substrate detection solution (Millipore, Watford, UK). Band quantification was determined using BioRad Image Lab Software.

### Immunoprecipitations

Muscle powder was homogenized, and protein quantified as above and 500µg protein was placed into a tube containing 25µL of antibody loaded Protein G-Dyna beads were aliquoted into an Eppendorf tube and prepped for immunoprecipitation using the instructed protocol (Thermo Scientific, Protein G-Dyna Beads). Antibodies for pulldown were used at a concentration of 1:100, and the final solution was submerged in a 30µL of 1X LSB, boiled for 5 minutes and stored at −80°C. 6µL of sample was loaded per well onto a Criterion TGX Stain-Free Precast Gel and carried forward with the western blotting protocol above.

### Antibodies

Primary antibodies for western blotting and immunoprecipitation were diluted to a concentration of 1:1000. Antibodies were from Cell Signaling Technology (Danvers, MA, United States)-total eEF2 (CS-2332S), p-53 (CS-2524S), phospho-eEF2 (CS-2331S), SIRT1 (CS-947S), Ac-Lys (CS9441S), phospho-S6 (CS-5364S), Ac-p53 (CS-252S), P-AKT (Ser473) (CS-4060S), Cytochrome-C (CS-4280S); Santa Cruz Biotechnology (Santa Cruz, CA, United States)-rps6 (SC-13007), rpL13a (SC-390131), Dystrophin (SC-465954); Abcam (Cambridge, UK)-Total OxPhos (ab110413); and Millipore-puromycin (MABE343).

### Statistics

Data were analyzed using one-way or two-way ANOVA (loading x inhibition) using GraphPad Prism software (GraphPad Software, Inc., La Jolla, CA). Tukey’s post hoc analysis was used to determine differences when interactions existed. Statistical significance was set at p < 0.05. All data are presented as mean ± standard error mean (SEM).

## Results

### SIRT1 Inhibitor Screen

The high-throughput screen of 800 natural products identified 45 compounds that inhibited SIRT1 >65% at a concentration of 50µM. Of these, many showed a dose-dependent effect on SIRT1, with 35 compounds showing at least 20% inhibition at 5µM (data not shown). Of the 35 inhibitors identified, 3 were already used extensively in human foods (Table 1) and came from 3 distinct chemical classes (one each quinone-methide, polyphenol, and flavonoid). These compounds (Celastrol, Epigallocatechin-3-Monogallate, and Epicatechin Monogallate) were selected to study effects on adaptive muscle hypertrophy in rats.

**Table 1.** Percent inhibition of SIRT1 deacetylase activity for the 3 compounds used in current study

| Compound Name | 5μM | 50μM | Class |
| --- | --- | --- | --- |
| Celastrol | 101 | 100 | Quinone-Methide |
| Epigallocatechin-3-Monogallate | 72 | 99 | Polyphenol |
| Epicatechin Monogallate | 66 | 99 | Flavinoid |

List of specific inhibitors of SIRT1 and the percent inhibition at 5 or 50 μM

### DOE Model Generation

The three compounds identified above were entered into a Box-Behnken incomplete factorial design to quickly assess any interactions between the different products. Thirteen rats received different combinations of the three products (ranging from 0-2mg/kg/day epicatechin; 0-10 mg/kg/day epigallocatechin-3-gallate; and 0-500µg/kg/day celastrol), while five controls received the middle amount of each product (1mg/kg/day epicatechin; 5mg/kg/day epigallocatechin-3-gallate; and 250µg/kg/day celastrol)to determine biological variability (Supplemental Table 1). Fourteen days of overload resulted in changes in muscle fiber cross-sectional area that ranged from −4.3% to 115.8% depending on the treatment (Figure 1), with the controls averaging 66.8±6.9%. From these data, response surface plots indicated that the epigallocatechin-3-gallate could modulate the effect of the other two products and a model predicting the optimal combination and concentration of each product was produced (Figure 1B).

**Figure 1.**
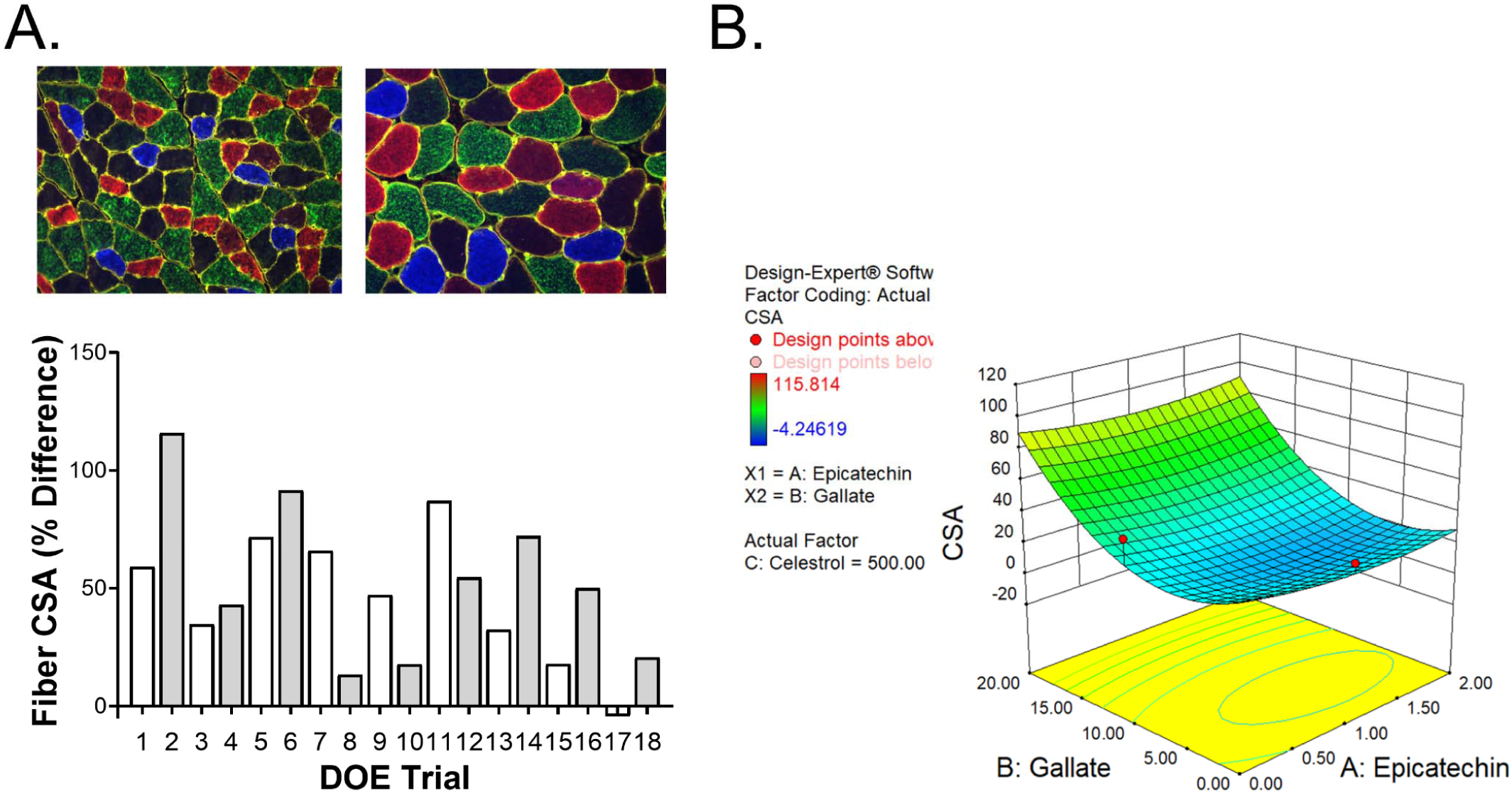
Development of a Box-Behnken model of natural products versus muscle cross-sectional area. Eighteen separate combinations and concentrations of the three natural products were established using an incomplete factorial design. Thirteen of the combinations were unique, whereas five were identical in order to determine the biological variability in the system. (A) Mean change in fiber cross-sectional area for each of the 18 different treatment groups following 14 days of overload. (B) Response surface plot for the relationship between CSA and the amount of epicatechin and epigallocatechin-3-gallate at a constant level of celastrol (500µg/kg/day).

### Model Validation

To validate the model, an independent group of rats underwent synergist ablation and were gavaged daily (beginning 5pm the night of surgery) with either a saline control or the predicted optimal, least effective, or a combination of the three products predicted to produce an increase in fiber CSA between the other two groups. The dose of each product for each group is outlined in Table 2. Following 14 days of overload and treatment, the animals were sacrificed, and body, heart, liver and muscle weights were determined (Figure 2). There were no statistical differences between the untreated and treated groups for body, heart, or liver mass, suggesting that the treatment did not result in any acute toxicity. Both the moderate and optimal groups showed a significant increase in muscle mass relative to the control treated rats. Analysis of muscle fiber CSA demonstrated that the control legs showed similar distribution of fiber CSA regardless of treatment. There was a right shift in fiber CSA with overload that was greatest in the optimal group. The mean fiber CSA of the SHAM legs for all groups were 1847±114.6, 1945±132.5, 1883±114.6, and 1730±60.0 µm^2^ for the control, least, moderate, and optimal groups, respectively, whereas the overloaded legs showed averages of 1901±108.3, 2348.1±172.8, 2306.5±119.7, and 2800±145.9 µm^2^, for the respective groups (Figure 2G). To test the predictive power of the Box-Behnken model, the predicted change in fiber CSA was plotted against the measured value for each of the groups. The resulting line had an r^2^ value of 0.9586 validating the ability of the model to predict changes in muscle hypertrophy (Figure 2H).

**Table 2.** Inhibitor compounds and the dose used during the validation study

| Compound Name | Optimal Dose | Average Dose | Least Dose |
| --- | --- | --- | --- |
| Celastrol | 0.5 | 0.2 | 0.435 |
| Epigallocatechin-3-Monogallate | 20 | 14 | 6 |
| Epicatechin Monogallate | 0.7 | 0.7 | 1.3 |

Doses of epigallocatechin-3-monogallate and epicatechin monogallate (mg/kg/day) and celastrol ( $\mu\text{g/kg/day}$ ) for each inhibitor cocktail used in the validation study.

**Figure 2.**
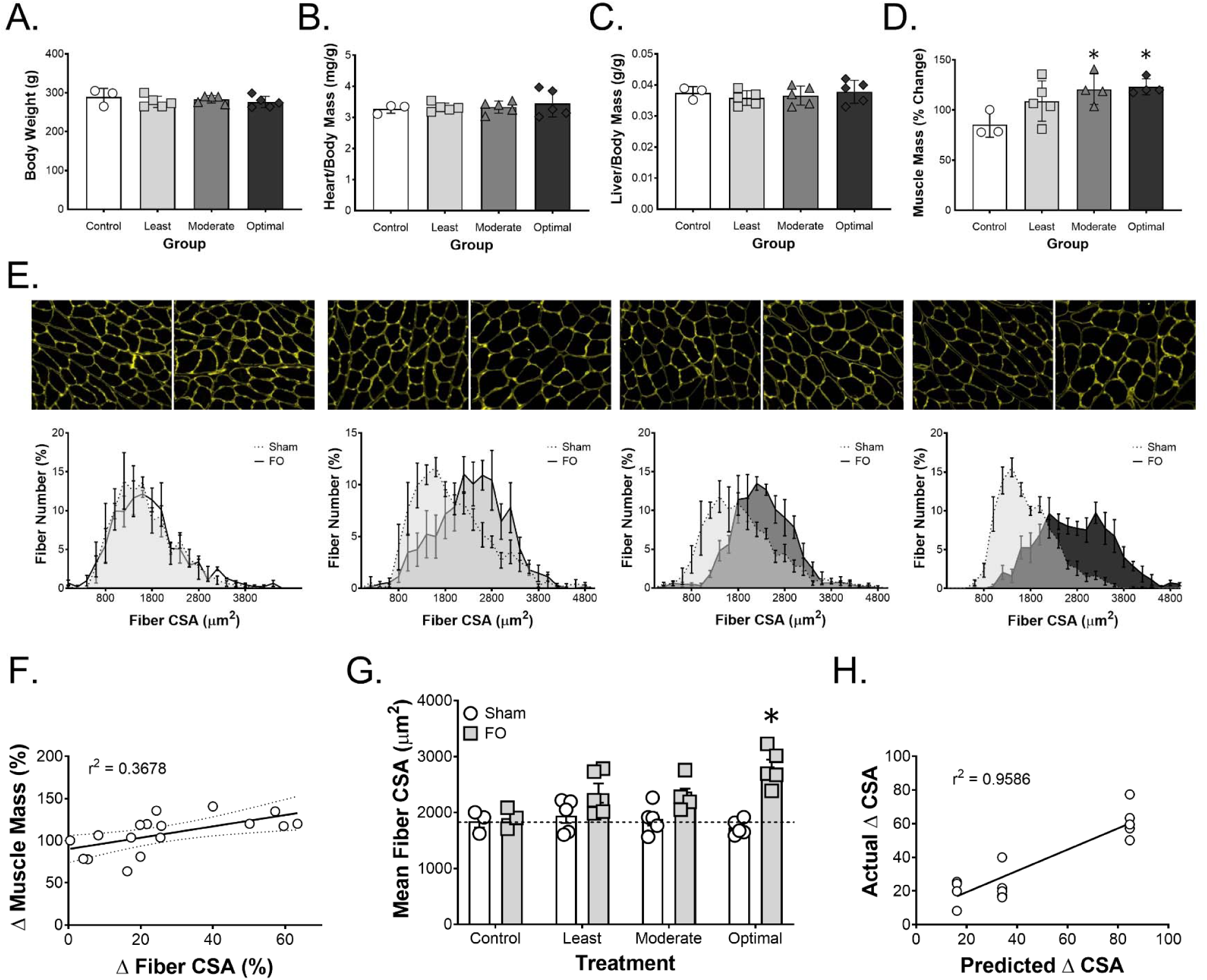
Validation of the Box-Behnken model of the relationship between natural products and muscle fiber cross-sectional area. (A) Body weight, (B) heart weight/body weight, (C) liver weight/body weight, (D) muscle mass following 14 days of overload with different levels of natural products. (E) Distributions of fiber cross-sectional area changes in the control, least effective, moderately effective, and most effective combination of natural products, from left to right. (F) Relationship between muscle fiber CSA and muscle mass following 14 days of overload. Note that mass increases ∼80% before a measurable change in mean fiber CSA occurs. (G) Mean fiber CSA as a function of overload and treatment. (H) Relationship between the prediction of the change in fiber CSA from the Box-Behnken model and the actual measured change in fiber CSA following 14 days of overload. Data are means ± SEM for n = 5 animals per treatment group. * indicates a significant difference from control muscle.

### SIRT1 Levels and Activity

Since the treatment was meant to inhibit SIRT1, the levels of SIRT1 and a gauge of its enzyme activity (p53 acetylation) was determined (Figure 3). As observed previously (13), the levels of SIRT1 increased significantly following overload in the control group (∼2-fold) and SIRT1 levels were even higher in both the control and overloaded limbs following treatment with the SIRT1 inhibitors. As a measure of SIRT1 activity, we determined the levels of acetylation of p53 at lysine 382. As has been reported for other SIRT1 inhibitors (25), overload together with treatment with the natural product cocktails increased p53 acetylation at this residue; however, there was no difference in p53 acetylation with the different doses. Lastly, to determine whether the natural product cocktail altered global protein acetylation, total acetylated proteins were measured and there was no statistical difference in total acetylated proteins with any of the treatments at the two-week time point.

**Figure 3.**
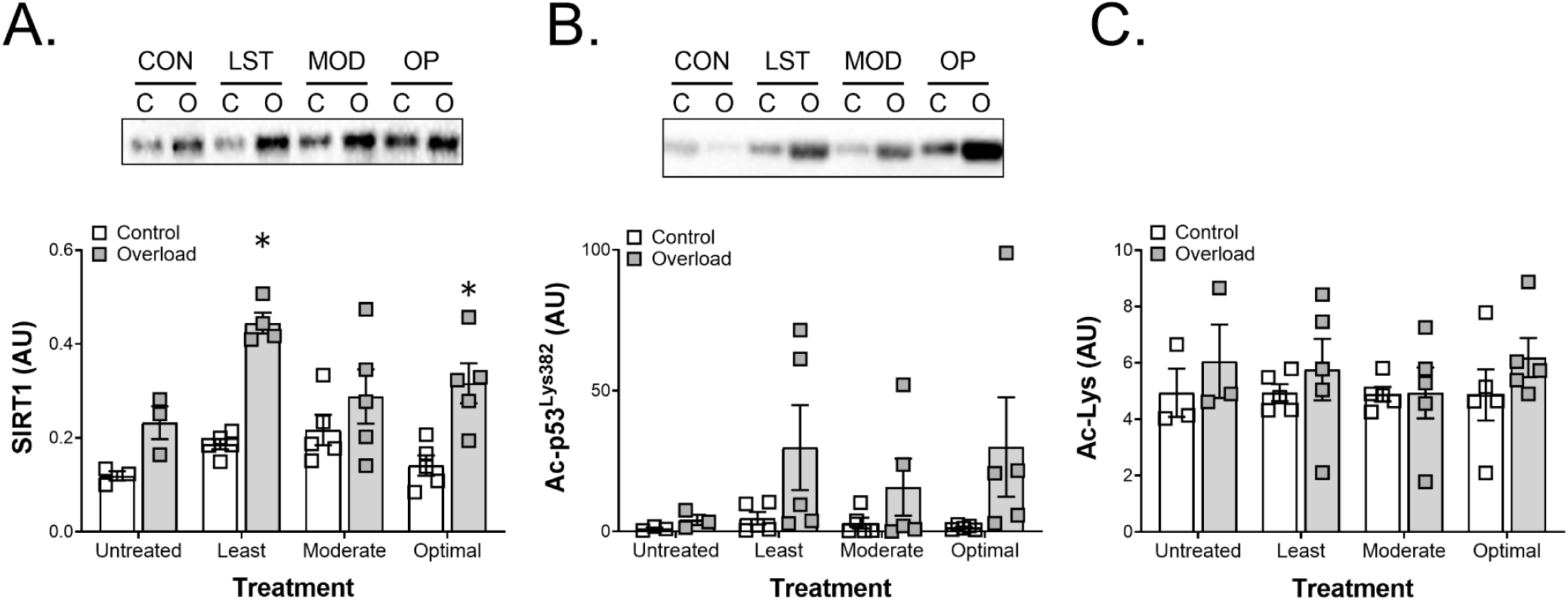
SIRT1 levels and acetylation with overload and natural product treatment. (A) Levels of SIRT1 protein following overload and natural product treatment. (B) Acetylation of p53 at K382 with both overload and treatment with natural products. Note the greater increase in p53 acetylation with the natural product treatment. However, the variability in each group precludes statistical significance. Finally, (C) shows the level of acetylation of lysines in the whole muscle homogenate. Data are means ± SEM for n = 5 animals per treatment group with every point shown. * indicates a significant difference from control muscle.

### Protein Synthetic Response

To begin to understand the mechanism through which the natural products were increasing muscle fiber CSA, the rate of protein synthesis was determined by SuNSeT. Even though there was a trend for baseline protein synthesis to increase with the natural product cocktail, the increase in protein synthesis with overload was similar across all treatment groups (Figure 4A). Since ribosome mass is thought to control protein synthesis in extreme states, such as during overload (11), we next determined total RNA and the rate of ribosome biogenesis. Contrary to our hypothesis, total RNA tended to decrease from control to optimal treatment. Further, when the rate of ribosomal RNA synthesis was determined by measuring the expression of the internal transcribed spacer 1 (ITS1) and 5⍰ external transcribed spacer (5⍰ETS), the expression of these markers of ribosomal biogenesis decreased from control, to least, to moderate, to optimal where the 5’ETS value was significantly lower than the control treated muscles (Figure 4C and D). To determine whether the increased growth in the natural product groups was the result of greater Akt-mTORC1 signaling, the phosphorylation of Akt, S6K1 and eEF2 were determined. There was a tendency for Akt phosphorylation to increase with overload and decrease in the natural products (Figure 4E); however, neither of these effects reached significance. As reported previously, S6K1 phosphorylation was higher in the overloaded leg (Figure 4F). Contrary to expectation, there was a trend for overload-induced S6K1 phosphorylation to decrease from control towards the optimal natural product combination; however, the activity of S6K1 (determined through eEF2 phosphorylation) was no different in any of the overloaded groups (Figure 4G).

**Figure 4.**
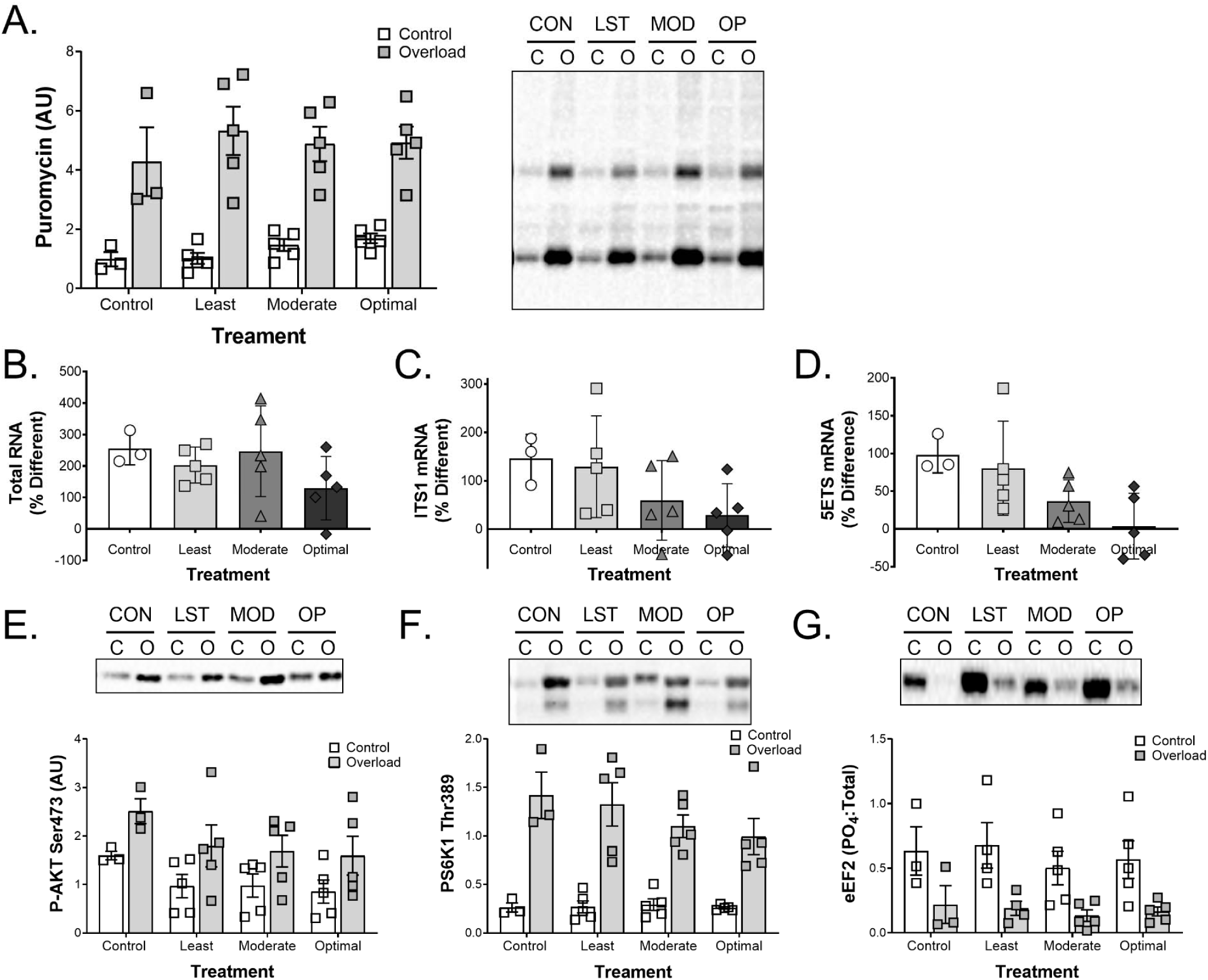
Protein synthesis and ribosomal markers with overload and natural product treatment. (A) Protein synthesis as estimated by SUnSET. Both the blot and quantified data are shown. Estimates of ribosomal mass were made by measuring (B) total RNA (∼80% of which is ribosomal RNA), (B) the internal transcribed spacer 1 (ITS1), and (C) the 5⍰ external transcribed spacer (5⍰ETS). To get an idea of Akt-mTORC1 signaling, (D) Akt Ser473, (E) S6K1 Thr389, and (F) eEF2 phosphorylation were measured. Data are means ± SEM for n = 5 animals per treatment group with every point shown. * indicates a significant difference from control muscle.

### Markers of Protein Turnover/Degradation

Since there was no effect of the natural products on protein synthesis, a quick measure of markers of protein turnover was made by measuring the expression of MuRF and MafBx. As has been reported previously, MuRF and MafBx expression tended to increase with overload and were not affected by the natural product treatment (Figure 5).

**Figure 5.**
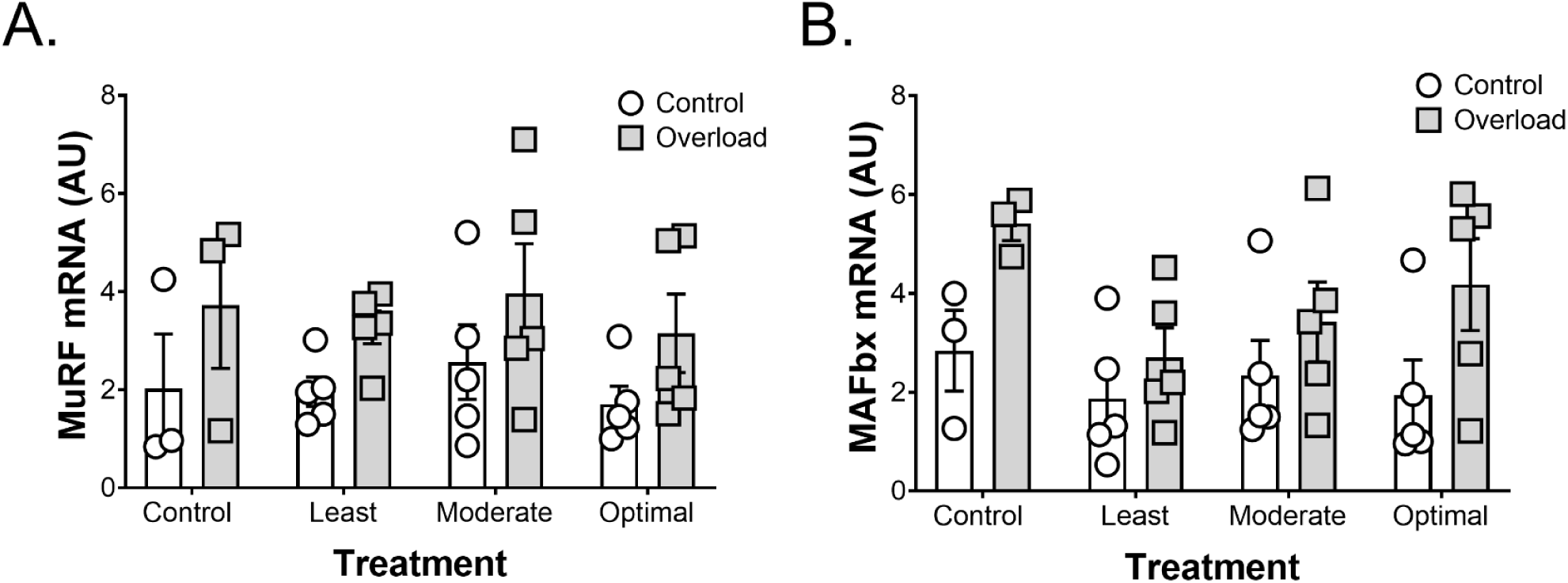
Markers of protein turnover with overload and natural product treatment. (A) MuRF and (B) MaFBx mRNA were measured. Data are means ± SEM for n = 5 animals per treatment group with every point shown. * indicates a significant difference from control muscle.

### Ribosomal Protein Acetylation

The ribosomal proteins are regulated by acetylation. Since SIRT1 is a deacetylase, the acetylation of proteins representative of the small and large ribosomal subunits was determined following immunoprecipitation. With optimization of the natural product cocktail, there was a trend for the acetylation of the ribosomal proteins to increase (Figure 6A and B). By contrast, S6K1 acetylation tended to decrease with optimization of the natural product cocktail (Figure 6C).

**Figure 6.**
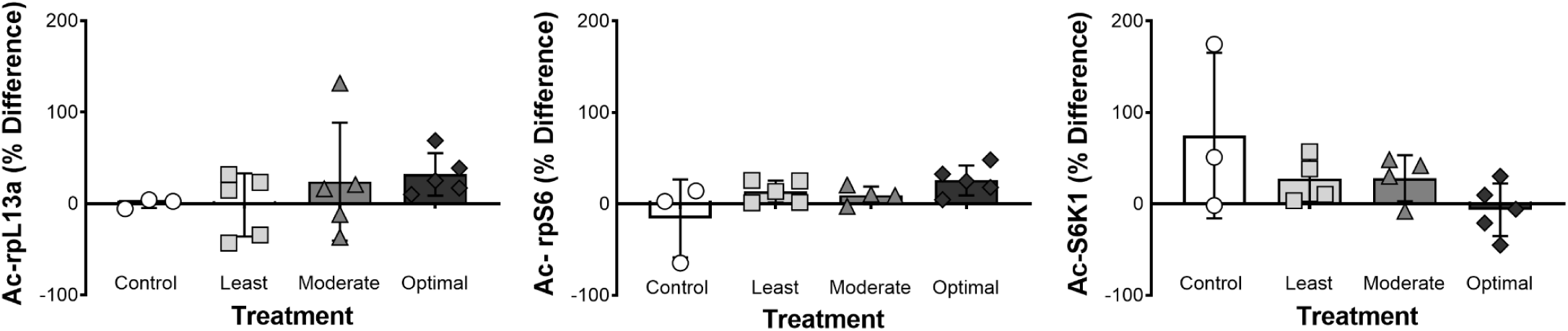
Acetylation of ribosomal proteins and regulators with overload and natural product treatment. To get an estimate of ribosomal acetylation, representative proteins from the large (A) L13A and small (S6) ribosomal subunits were blotted following immunoprecipitation with an acetyl-lysine antibody. (C) Levels of S6K1 acetylation were also determined in the same manner. Note the opposite pattern of the two measures. Data are means ± SEM for n = 5 animals per treatment group with every point shown. * indicates a significant difference from control muscle.

## Discussion

Here we show that several natural products have the ability to inhibit SIRT1 in an *in vitro* activity assay. Combining three of these natural products that are generally recognized as safe (GRAS), a Food and Drug Administration designation that a chemical added to food is considered safe by experts, and therefore exempt from the Federal Food, Drug, and Cosmetic Act food additive tolerance requirements, in the optimal manner results in a significant increase in muscle fiber hypertrophy following 14 days of overload. The significant increase in muscle fiber CSA was not the result of an increase in ribosomal mass. In fact, the optimal group showed significantly less 5’ETS and a strong trend towards lower ITS1 levels and total RNA, suggesting fewer ribosomes than the control muscles. The acetylation of ribosomal proteins tended to increase suggesting that the increase in myofibrillar protein could be the result of an increase in ribosomal efficiency rather than capacity. Importantly, the natural product cocktail did not alter body mass or the weight of the heart and liver suggesting that it has limited toxicity and together with exercise may be useful in growing or maintaining muscle mass.

We have previously identified SIRT1 as one of a series of molecular breaks that limit the growth of muscle in response to extreme stimuli, such as synergist ablation (13). We hypothesized that the activation of SIRT1 would result in the deacetylation of TAF68, a component of the SL-1 transcription factor that drives the expression of the 47S ribosomal RNA (20). Deacetylation of TAF68 has previously been shown to inhibit rRNA transcription and therefore ribosome mass. Since ribosome mass is thought to limit growth following synergist ablation (11, 17), we hypothesized that blocking SIRT1 would decrease TAF68 acetylation, increase the expression of rRNA, increase the capacity for protein synthesis, and allow greater skeletal muscle hypertrophy in response to overload. With this background, we sought to determine whether SIRT1 could be inhibited by natural products and produce an improvement in growth.

Using the NatProd Collection, 800 pure natural products and their derivatives, derived from plant, animal, and microbial sources, we identified 45 compounds that inhibited the activity of SIRT1 towards p53 by greater than 65% at a concentration of 50µM and 35 compounds that inhibited SIRT1 by at least 20% inhibition at 5µM. This represents a unique list of compounds, many of which are polyphenols, including quinones and flavonoids that inhibit SIRT1. The fact that the majority of the compounds that inhibited SIRT1 were polyphenols suggests that regulation of SIRT1 may be one reason that polyphenols have a significant impact on human health and disease prevention (22). We chose to focus on three of these polyphenols, epicatechin, epigallocatechin-3-gallate, and celastrol because they had a history of use in human medical trials without complication (14, 28, 29) and have different chemical structures that might mean different degrees of digestion, absorption, delivery, and activity in muscle following ingestion.

Using an incomplete factorial design, we treated animals with different concentrations and combinations of natural products to create a model as to how each natural product contributes to the increase in muscle CSA following overload. Response surface plots were used to determine the relative importance of each component and their interaction with the other natural products in the cocktail (Figure 1B). To validate the model, we chose three different combinations of the natural products based on their predicted effect on muscle fiber CSA following overload. An independent cohort of rats (n=5 per treatment) underwent synergist ablation and then received a daily gavage containing of one of the three combinations of natural products, or the placebo control. The fact that the model prediction for the increase in fiber CSA was proportional to the measured change in CSA (r^2^ = 0.9586), suggests that the model was valid and that the predicted combination of epicatechin, epigallocatechin-3-gallate, and celastrol was optimal for muscle hypertrophy.

The optimal combination of epicatechin, epigallocatechin-3-gallate, and celastrol increased mean fiber CSA 61.5% compared with ∼4% in the control group. This finding is striking for two reasons. First, as seen in Figure 2F, the increase in muscle mass following overload was not proportional to the mean increase in fiber CSA. In fact, the mass of the muscle appeared to have to increase by ∼80% before an increase in the mean fiber CSA was observed. However, we did not observe a significant increase in fiber number over that period (data not shown). These data suggest that the majority of the increase in muscle mass that occurs following 14 days of functional overload is not due to an increase in average fiber CSA. In the plantaris muscle, there are regions of very big fibers and other regions of relatively small fibers. It is possible that the regional difference in fiber CSA negate any obvious effect on mean fiber area following overload. However, it is also possible that the muscle is growing in other ways. Following the removal of the soleus and gastrocnemius muscles, the ankle of the rat is held in a more dorsiflexed position, which would be expected to increase the resting length of the plantaris. We have preliminary data that indicates that one result of the shift in ankle position is that the plantaris muscle increases in length by approximately 10%. Others have recently made a similar suggestion in mice (16). These data suggest that the plantaris muscle is increasing in mass as a result of functional overload in part through the addition of sarcomeres in series. However, this hypothesis needs to be more thoroughly evaluated.

The finding that the optimal group had a 61.5% increase in mean fiber CSA compared with ∼4% in the control group is also striking for the magnitude of the difference in hypertrophy with the natural product cocktail. Other treatments that augment muscle hypertrophy, such as consuming leucine rich protein, have an effect size of ∼5% (3). These data suggest that the mechanism underlying the effect of the natural product cocktail is distinct and likely rate limiting in load-induced skeletal muscle hypertrophy. One proposed limit to skeletal muscle hypertrophy in both mice and man is the capacity for protein synthesis; i.e. ribosome mass (11, 17, 27). To determine whether ribosome mass was increased in the animals fed the natural product cocktail we determined total RNA relative to muscle mass. Contrary to our hypothesis, total RNA tended to increase less with the natural product cocktail than in the vehicle controls. In support of this observation, the rRNA spacers (ITS1 and 5’ETS) showed the same pattern, with the change in 5’ETS being statistically lower than controls. These data suggest that even though the natural product cocktail increased hypertrophy, the improvement was not the result of an increase in translational capacity. Alternatively, the natural product cocktail may have resulted in an early (3-7 days) increase in ribosome mass that returned to baseline sooner. Further experiments are necessary to determine whether this is a difference in dynamics or absolute change in total RNA.

One possible explanation for the apparent increase in muscle protein without a concomitant rise in ribosome mass is an increase in translational efficiency. While there is a paucity of recent work on the role of acetylation of the ribosomal proteins and translational efficiency, early work showed that following hepatectomy, when protein synthesis rates increase to regenerate the tissue, the acetylation of the ribosomal proteins precedes the protein synthetic response (18). This suggests that acetylation of the ribosomal proteins may increase translational efficiency. Choudhary and colleagues identified 75 ribosomal proteins that were acetylated at a minimum of 136 locations (4). For the mitochondrial ribosome, proteins MRPL10 and 19 are deacetylated by the mitochondrial sirtuin SIRT3 (30). When SIRT3 is overexpressed, MRPL10 and 19 become deacetylated and this corresponds with a decrease in protein synthesis. When a catalytically inactive SIRT3 is used there is no change in acetylation or protein synthesis. Further, when SIRT3 is targeted with shRNA, acetylation and protein synthesis both increase (30). Lastly, ribosomes isolated from the liver of SIRT3 knockout mice show more protein synthesis per unit of ribosomal protein (30). Together, these data suggest that sirtuins can deacetylate ribosomal proteins and this corresponds with a decrease in translational efficiency. Consistent with these data, we show that our putative SIRT1 inhibiting natural product cocktail tends to increase the acetylation of two ribosomal proteins and this is associated with a greater change in protein synthesis (similar increases in puromycin) relative to total RNA or ribosomal biogenesis (5’ETS), suggesting improved translational efficiency.

Beyond the acetylation of ribosomal proteins, the cell size regulator S6K1 can also be acetylated (9, 10) and this decreases its ability to be phosphorylated by mTORC1 (15), and its ability to phosphorylate ribosomal protein s6 in mesangial cells (6, 15). The deacetylation of S6K1 can be catalyzed by either SIRT1 or 2 (15). Therefore, inhibition of SIRT1 would be expected to increase S6K1 acetylation and decrease Thr389 phosphorylation. Interestingly, in muscles treated for 14 days with presumed inhibitors of SIRT1, we observed a tendency for S6K1 acetylation and phosphorylation to both decrease. When trying to rectify our data with the existing literature, it is important to note that the effect of SIRT1 on S6K1 acetylation was performed in culture following 3 hours of treatment with the sirtuin inhibitor nicotinamide (15). It is also important to note that much of the *in vitro* study looked specifically at S6K1 acetylation at lysines 427, 484, 485, and 493. By contrast, the current work looked at total acetylation of S6K1 following immunoprecipitation. It is possible that longer periods of sirtuin inhibition in the presence of a growth stress would lead to the acetylation of 427, 484, 485, and 493 but deacetylation at other sites, resulting in the net deacetylation that was observed in the current study. Further work is needed to better understand the specific residues acetylated in S6K1 and their role in the regulation of protein synthesis.

## Conclusions

Using an *in vitro* assay, we have identified several natural products that can inhibit SIRT1 activity. By combining three of these inhibitors *in vivo* we were able to develop a model for how different combinations contributed to muscle hypertrophy following overload. When validating the model, we discovered that the optimal combination of natural products could significantly increase muscle hypertrophy in response to loading even though it significantly decreased ribosome biogenesis and tended to decrease the rise in ribosome mass that occurs with overload. This suggests that the natural product described here may increase ribosome efficiency, possibly by increasing the acetylation of ribosomal proteins, resulting in greater skeletal muscle hypertrophy.

## Acknowledgements

The work was supported by a research grant by Advanced Muscle Technologies (KB) and a grant from the National Institute of Arthritis, Musculoskeletal and Skin Diseases (AR069775R21) to SS and KB.

## Conflict of Interest

The inhibition of SIRT1 and enhanced muscle hypertrophy (U.S. patent number 9,480,675) and the natural product cocktail described in this manuscript (U.S. patent number 11,312,695) have been patented. Both patents have been licensed by Advanced Muscle Technologies (AMT). AMT provided funds for the studies described in this manuscript, paid KB as a consultant, and provided shares to SS and KB.

## Figure Legends

**Supplemental Table 1.**
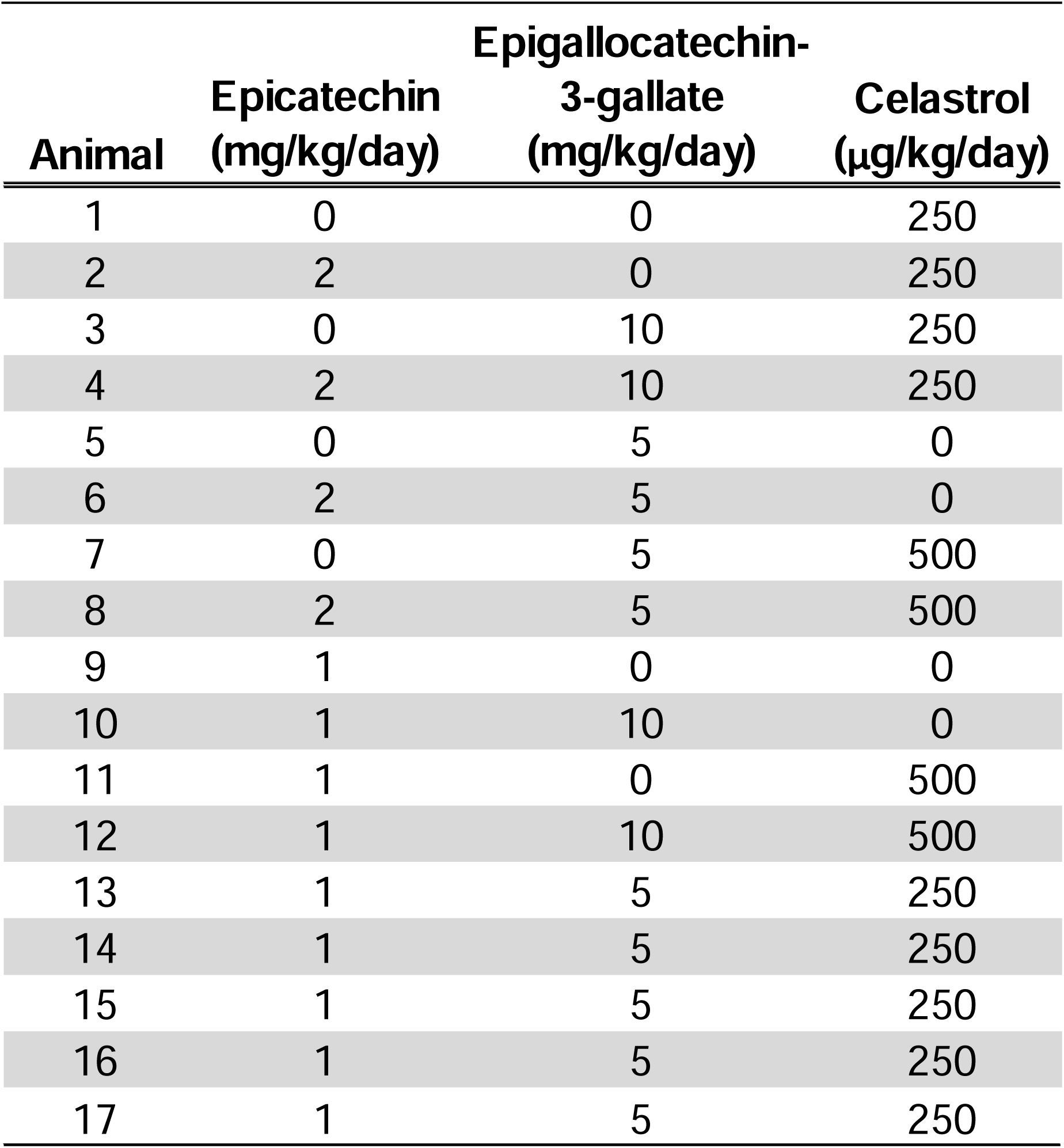
Amount of natural product given to each animal nightly for the DOE experiment.

